# Development and behavior of *Steinernema hermaphroditum* nematodes are impacted by a bacterial flavin monooxygenase

**DOI:** 10.64898/2026.07.20.739592

**Authors:** Tyler G. Myers, Heidi Goodrich-Blair, Jennifer K. Heppert

## Abstract

Beneficial animal-microbe symbioses occur across the tree of life, with observable impacts on animal development and behavior. Some of these impacts occur when microbial enzymes act upon host-derived metabolites. The bacterial enzyme complex HpaBC hydroxylates aromatic compounds, including tyrosine and precursors of the neurotransmitter dopamine, raising the possibility that this enzyme may indirectly impact neurotransmitter-dependent phenotypes. The bacterium *Xenorhabdus griffiniae*, a beneficial intestinal symbiont of *Steinernema hermaphroditum* entomopathogenic nematodes, encodes HpaBC. Computational docking of *X. griffiniae* HpaB affinity for dopamine pathway substrates revealed energetically favorable docking of both dopamine and tyrosine, though neither was as favorable as the predicted affinity of HpaB for the canonical substrate, 4-hydroxyphenylacetate. We hypothesized that *Xenorhabdus* HpaBC may influence nematode host development or behavior through its potential action on tyrosine degradation or dopamine catabolism. In support of this idea, we demonstrate a negative correlation between *X. griffiniae hpaBC* expression and body length and egg-laying behavior in adult *S. hermaphroditum* nematodes. Moreover, *hpaBC* expression is necessary for *X. griffiniae* bacteria to colonize the nematode intestine in the infective juvenile stage. Taken together, these findings indicate that HpaBC may be a part of the metabolic crosstalk occurring between *X. griffiniae* and *S. hermaphroditum* at multiple stages of their shared life cycle. *hpaBC* orthologs are widely present across bacterial phyla including in pathogens and mutualists of plants and animals. Our findings raise the possibility that HpaBC could be a conserved mechanism by which host-associated bacteria influence physiology and behavior.

**Importance:** Beneficial animal–microbe interactions can shape development, physiology, and behavior in organisms across the tree of life, yet in many cases, the molecular mechanisms driving these interactions remain poorly understood. This study reveals that the bacterial metabolic enzyme complex HpaBC plays a role in the *Steinernema–Xenorhabdus* symbiosis, influencing nematode growth, reproduction, and host colonization. As *hpaBC* expression is conserved across many bacterial phyla, these findings highlight a potentially widespread mechanism for modulating animal physiology through microbial metabolism.

## Introduction

Across diverse animal species, beneficial interactions with microbes regulate growth, developmental timing, tissue differentiation, and behavior (1–3). These impacts occur through the exchange of nutrients between host and microbe, and through the production or modification of bioactive signaling molecules, such as neurotransmitters. In the human intestine, microbes modify host neurotransmitter levels through direct synthesis or degradation, the creation of neurotransmitter precursors, and more indirectly through the production of metabolites that stimulate host neurotransmitter synthesis or release (4). Additionally, many studies have correlated the presence of specific microbial taxa with host behavioral and physiological changes (5–8). Despite these advances, dissecting the mechanistic impacts of microbes in humans is challenging because of the heterogeneity and variability of the animal intestinal microbiome, the diversity of intestinal cell types, connections with both the enteric and central nervous systems, and the complexities associated with measuring human behavior.

Laboratory animal models such as nematodes can be powerful platforms for mechanistic investigations of host-microbe interactions because of their relative simplicity and tractability. In the free-living nematode *Caenorhabditis elegans,* microbes alter multiple behaviors including feeding, motility, reproduction, and stress responses through both nutrient-dependent mechanisms and direct neurochemical modulation (9). A striking example is 2,5-diketopiperazine, a microbial metabolite gliotoxin which synergizes with endogenous nematode serotonin to acutely stimulate egg-laying behavior in hermaphrodites (10). Egg laying in *C. elegans* is orchestrated by the hermaphrodite-specific serotonergic HSN motor neurons, whose activity integrates excitatory serotonergic drive with inhibitory dopaminergic and cholinergic signaling (11–14). These neurons respond robustly to exogenous neurotransmitters and synthetic 5-HT receptor agonists, positioning the nematode egg-laying circuit as a sensitive readout of microbial influence on host neuromodulation (13). Microbial cues also guide nematode decision-making. *C. elegans* differentiates between nutritive and pathogenic bacteria through detection of specific microbial metabolites, including polyamines that activate defined sensory and intraneuronal pathways (15). Microbes provide essential nutrients such as heme, and variation in the composition of environmental bacterial communities can determine whether *C. elegans* populations undergo reproductive development or form dauers, an alternative, stress-resistant larval stage (16,17). Within the intestinal tract, neuroendocrine pathways can detect colonizing bacteria and changes in physiology leading to activation of an immune response, further illustrating the extensive integration of microbial signals into nematode host developmental and behavioral decision-making (18).

Complementary to studies using free-living *C. elegans* as a model system, entomopathogenic nematodes of the genus *Steinernema* can be used to investigate the reciprocal influence between hosts and symbionts on their respective physiologies. In contrast to *C. elegans*, whose intestinal microbiome consists of a consortium of species, *Steinernema* nematodes maintain evolutionarily stable, species-specific partnerships, or symbioses, with bacteria of the genus *Xenorhabdus* (19,20,17,21). The nematodes are colonized in the anterior intestine by their *Xenorhabdus* symbionts (20), and together this organismal pair infects and kills insects, using the cadaver as a primary nutrient source (22). When nutrients become limiting within the exhausted insect cadaver, the *Xenorhabdus* bacteria re-colonize stage-specific tissues within the nematode, ultimately populating a specialized compartment within the lumen of the anterior intestine called the receptacle in the nematode infective juvenile (IJ) stage, where they are vectored by the nematode to a new insect host (23). The close symbiotic relationship between *Steinernema* nematodes and *Xenorhabdus* bacteria offers a distinctive opportunity to explore how bacterial products influence host behavior.

One *Xenorhabdus* metabolic pathway with the potential for perpetuating host influence is the HpaBC enzyme complex, a two-subunit monooxygenase system involved in downstream tyrosine catabolism across bacterial phyla (24–26). HpaB is an FADH₂-dependent monooxygenase and HpaC is a flavin reductase that regenerates reduced FADH₂ following each catalytic cycle (27). Beyond its canonical substrate 4-hydroxyphenylacetate (4-HPA), HpaBC displays notable substrate promiscuity, acting on diverse phenolic and catecholic compounds (28,29). Engineered variants of HpaBC can synthesize dopamine *in vitro* from L-tyrosine (28), raising the possibility that naturally-occurring bacterial HpaBC activity may affect animal neuromodulatory pathways. More recently, clinical applications harnessing the influence of HpaBC on dopamine synthesis have been explored as a gut-based treatment for Parkinson’s disease (30). To better understand the interplay between bacterial HpaBC and animal behaviors, we used the *hpaBC-*encoding bacterium, *Xenorhabdus griffiniae* HGB2511 and its nematode host, *Steinernema hermaphroditum* (31,32). Building on preliminary observations suggesting *Xenorhabdus hpaBC* involvement in nematode host-associated phenotypes and changes in neurotransmitter levels throughout the life cycle (22,33), we hypothesized that altering *X. griffiniae hpaBC* expression could influence both developmental and behavioral outcomes in the nematode host. We found that elevated *hpaBC* expression correlates with reduced adult body length and suppressed egg-laying behavior in *S. hermaphroditum*. We further demonstrate that *hpaBC* activity is required for bacterial colonization of the receptacle in the IJ nematode, making it necessary for bacterial transmission in subsequent generations. Together, these findings uncover a previously unappreciated role for HpaBC in shaping nematode development, reproduction, and host colonization, expanding our understanding of how bacterial enzymatic activity drives host phenotypic traits within nematode–microbe symbioses.

## Materials and Methods

### Plotting substrate docking to HpaB

To compare the predicted binding affinities of HpaB enzymes to canonical (*i.e.,* 4-HPA) and non-canonical (*i.e.,* dopamine and L-tyrosine) substrates, the HpaB subunit amino acid sequences and three-dimensional structures were retrieved from the InterPro database or the MaGe MicroScope platform from *Photorhabdus luminescens* and eighteen *Xenorhabdus* strains (https://mage.genoscope.cns.fr/microscope/) (34–36) (Table 1). Corresponding three-dimensional protein structures were obtained as PDB-format coordinate files, either directly from available structural predictions or by exporting predicted structures associated with these entries. For HpaB proteins from two strains, *X. griffiniae* HGB2511 and *X. nematophila* Websteri, structures were obtained in .cif format through MaGe Microscope and converted to .pdb format to ensure compatibility with downstream docking analyses. The canonical Simplified Molecular Input Line Entry System (SMILES) strings for 4-HPA (C1=CC(=CC=C1CC(=O)O)O), dopamine (C1=CC(=C(C=C1CCN)O)O), and L-tyrosine (C1=CC(=CC=C1C[C@@H](C(=O)O)N)O) were retrieved from PubChem (https://pubchem.ncbi.nlm.nih.gov) and provided as ligand inputs for SwissDock 2024 (37,38). Docking runs were performed for the selected ligands against each tested HpaB structure using the default blind-docking protocol in AutoDock Vina with a recording volume of 25,000 Å_3_, search space centered at [0,0,0], and sampling exhaustivity set to 4. The resulting predicted docking affinities were extracted from the SwissDock output. Finally, all affinity values for 4-HPA, dopamine, and tyrosine across the nineteen strains were imported into GraphPad PRISM for statistical comparison.

**Table 1.**
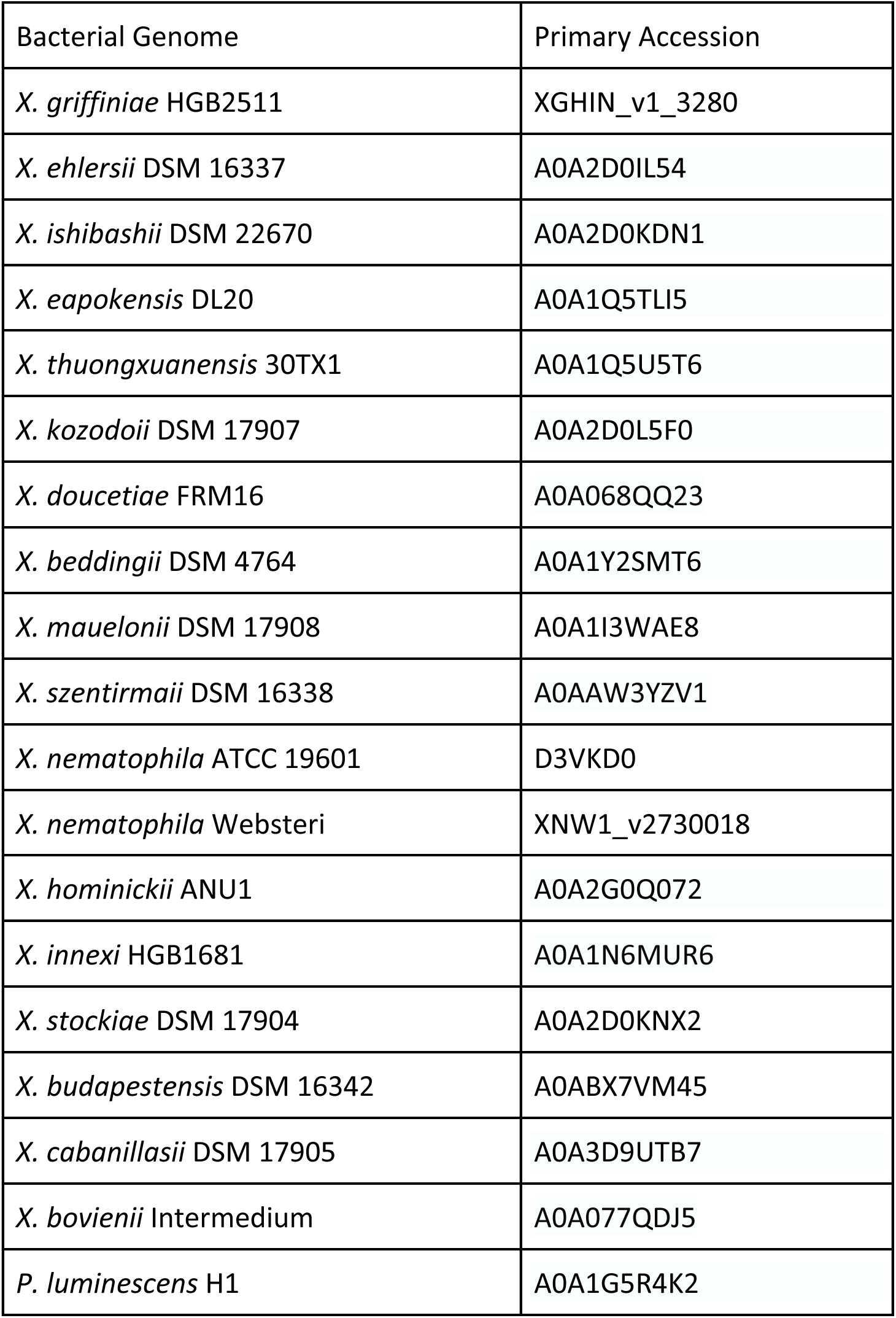
HpaB proteins assessed for predicted docking affinity.

| Bacterial Genome | Primary Accession |
| --- | --- |
| *X. griffinae* HGB2511 | XGHIN_v1_3280 |
| *X. ehlersii* DSM 16337 | A0A2D0IL54 |
| *X. ishibashii* DSM 22670 | A0A2D0KDN1 |
| *X. eapokensis* DL20 | A0A1Q5TLI5 |
| *X. thuongxuanensis* 30TX1 | A0A1Q5U5T6 |
| *X. kozodoii* DSM 17907 | A0A2D0L5F0 |
| *X. doucetiae* FRM16 | A0A068QQ23 |
| *X. beddingii* DSM 4764 | A0A1Y2SMT6 |
| *X. mauelonii* DSM 17908 | A0A1I3WAE8 |
| *X. szentirmaii* DSM 16338 | A0AAW3YZV1 |
| *X. nematophila* ATCC 19601 | D3VKD0 |
| *X. nematophila* Websteri | XNW1_v2730018 |
| *X. hominickii* ANU1 | A0A2G0Q072 |
| *X. innexi* HGB1681 | A0A1N6MUR6 |
| *X. stockiae* DSM 17904 | A0A2D0KNX2 |
| *X. budapestensis* DSM 16342 | A0ABX7VM45 |
| *X. cabanillasii* DSM 17905 | A0A3D9UTB7 |
| *X. bovienii* Intermedium | A0A077QDJ5 |
| *P. luminescens* H1 | A0A1G5R4K2 |

### Identification of *X. griffiniae* HGB2511 and *S. hermaphroditum* homologs in the dopamine metabolism pathway

Enzymes that mediate key steps in the tyrosine and dopamine metabolism pathways were first identified in the Kyoto Encyclopedia of Genes and Genomes (KEGG) database (39). Orthologs of these enzymes were identified in the *X. griffiniae* HGB2511 genome (CP147738.1) by a combination of genome annotation and BLASTp using the MaGe Microscope platform. *C. elegans* orthologs were identified using the KEGG Orthology annotations, literature, and BLASTp (40). *S. hermaphroditum* orthologs were inferred using a Biomart comparison of the *C. elegans* N2 (PRJNA13758) and *S. hermaphroditum* PS9179 (PRJNA982879) genomes and by Reciprocal Best BLAST, using BLASTp of the *C. elegans* genes previously identified. Orthologs were mapped onto a simplified version of the KEGG pathways (Fig. 1A), and enzyme names, accession, locus tag, and amino acid sequences identified can be found in Table S1.

**Figure 1.**
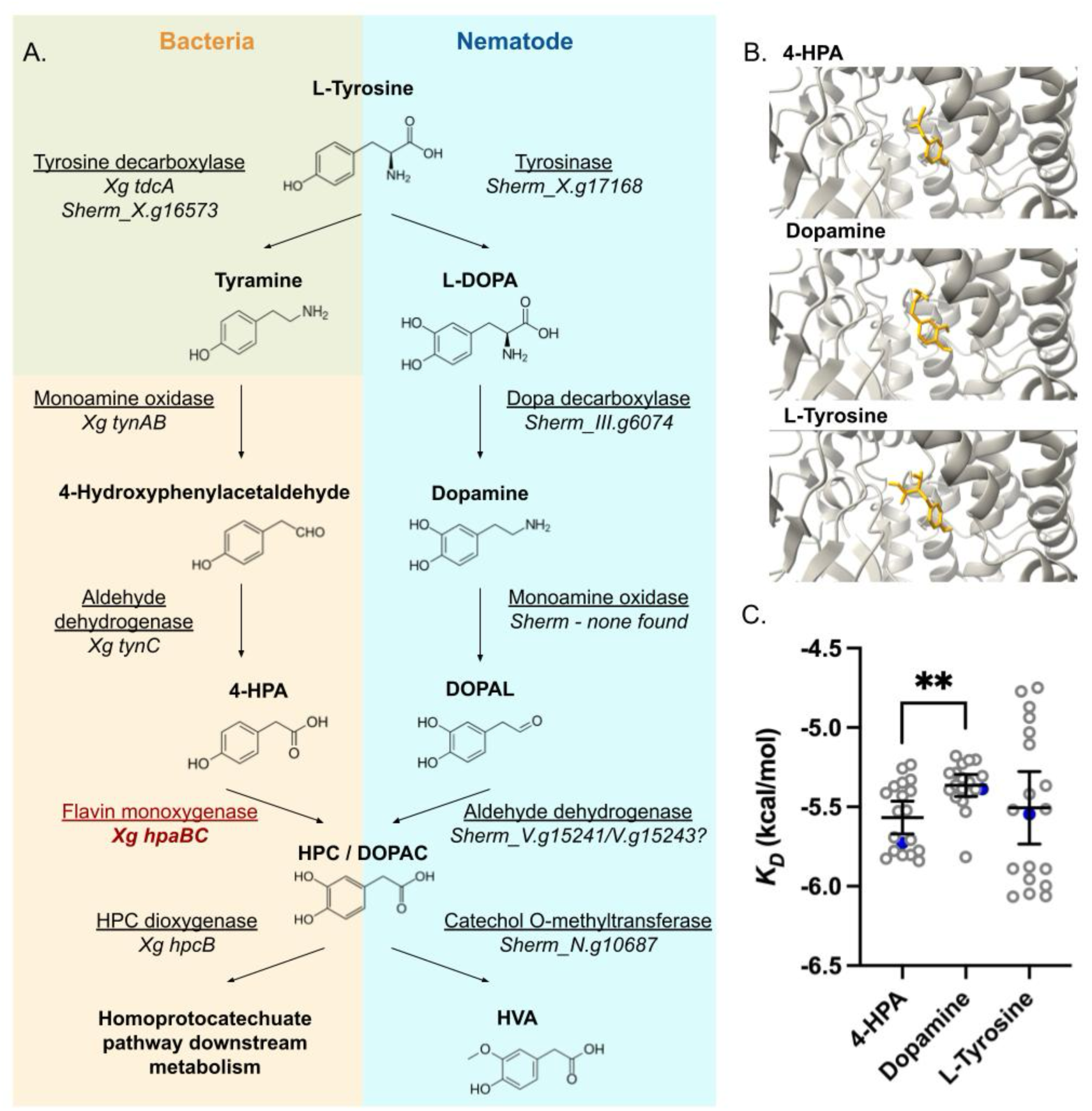
The bacterial HpaB enzyme may bind multiple substrates in the dopamine/tyrosine metabolic pathway based on *in silico* docking predictions. (A) The tyrosine and dopamine metabolism pathways in bacteria (tan) and nematodes (blue) and their overlap are depicted. (B) As visualized within SwissDock, the substrate molecules 4-HPA, dopamine, and L-tyrosine (yellow) occupy similar orientations within the active site of the predicted *X. griffiniae* HGB2511 HpaB enzyme (grey ribbon diagram). (C) 4-HPA, dopamine and L-tyrosine are each predicted to have energetically favorable affinities when docking within the active site of HpaB. The predicted docking affinity of HpaB for 4-HPA is significantly higher than that for dopamine (**, *P* < 0.01, two-way ANOVA). The predicted docking affinity of HpaB for L-tyrosine was not significantly different from either 4-HPA or dopamine. Blue dots represent the *K_D_* values for *X. griffiniae* HGB2511 HpaB, and error bars depict a 95% confidence interval.

### Generation of a *X. griffiniae* HGB2511 strain with arabinose-inducible HpaBC

Primers (Xg_hpaB_F and Xg_hpaB_500_R, Table 2) flanking a 500-bp region beginning at the predicted start codon of the *X. griffiniae* HGB2511 *hpaB* coding sequence (downloaded from the MaGe Microscope platform) were designed by eye using ApE (https://jorgensen.biology.utah.edu/wayned/ape/) (41) and used to amplify the predicted *hpaB* promoter fragment from *X. griffiniae* genomic DNA template through polymerase chain reaction (PCR). Ex Taq Polymerase, dNTPs, and buffer (TaKaRa Bio) were used according to the manufacturer’s instructions with a primer melting temperature of 61°C and completion of 30 cycles of amplification. The resulting amplicon was directionally cloned into the pCEP_kan plasmid backbone (Table 3) (42) immediately downstream of the *P_araBAD_* (*P_ara_*) promoter using NEBuilder HiFi DNA Assembly (New England Biolabs) according to manufacturer’s instructions. The resulting *P_ara_*::*hpaB*_500 construct was introduced into DAP-dependent *E. coli* S17 *λpir* (HGB1261, Table 3) by electroporation (MicroPulser Bio-Rad) resulting in the plasmid donor strain HGB2621. For conjugation, overnight cultures of HGB2621 and the *X. griffiniae* HGB2511 recipient strain were subcultured in a 1:10 ratio into fresh, antibiotic-free dark LB (liquid Luria-Burtani (LB) medium stored in the dark) (43) and grown at 30°C with agitation for 3 h until an OD_600_ of 0.8 was reached. A conjugation mixture was prepared by combining 900 µL of *X. griffiniae* with 100 µL of *E. coli* in a microcentrifuge tube and spinning at max speed for 1 min. The concentrated cell suspension was then spotted onto LBP (LB + 0.1% pyruvate) + 0.3 mM diaminopimelic acid (DAP) agar and incubated for 24 h at 30°C. The spot was scraped into sterile LB, and dilutions were plated onto LB + 50 µg/mL kanamycin (without DAP to prevent growth of the DAP-requiring donor *E. coli* strain). Individual kanamycin-resistant exconjugant colonies were transferred to a grid plate on LB + 50 µg/mL kanamycin and each was then re-streaked to obtain isolated clones prior to phenotypic and PCR verification of promoter insertion. Candidate exconjugants were initially confirmed to be *X. griffiniae* and not *E. coli* based on a catalase negative phenotype (44). For candidate exconjugants, the insertion of the *P_ara_* promoter upstream of *hpaB* was verified by comparing PCR amplification fragment sizes surrounding the *hpaBC* locus in HGB2622 (5.4 kb) and HGB2511 (0.5 kb) using Xg_hpaB_F and Xg_hpaB_R_1 primers (Table 2).

**Table 2.**
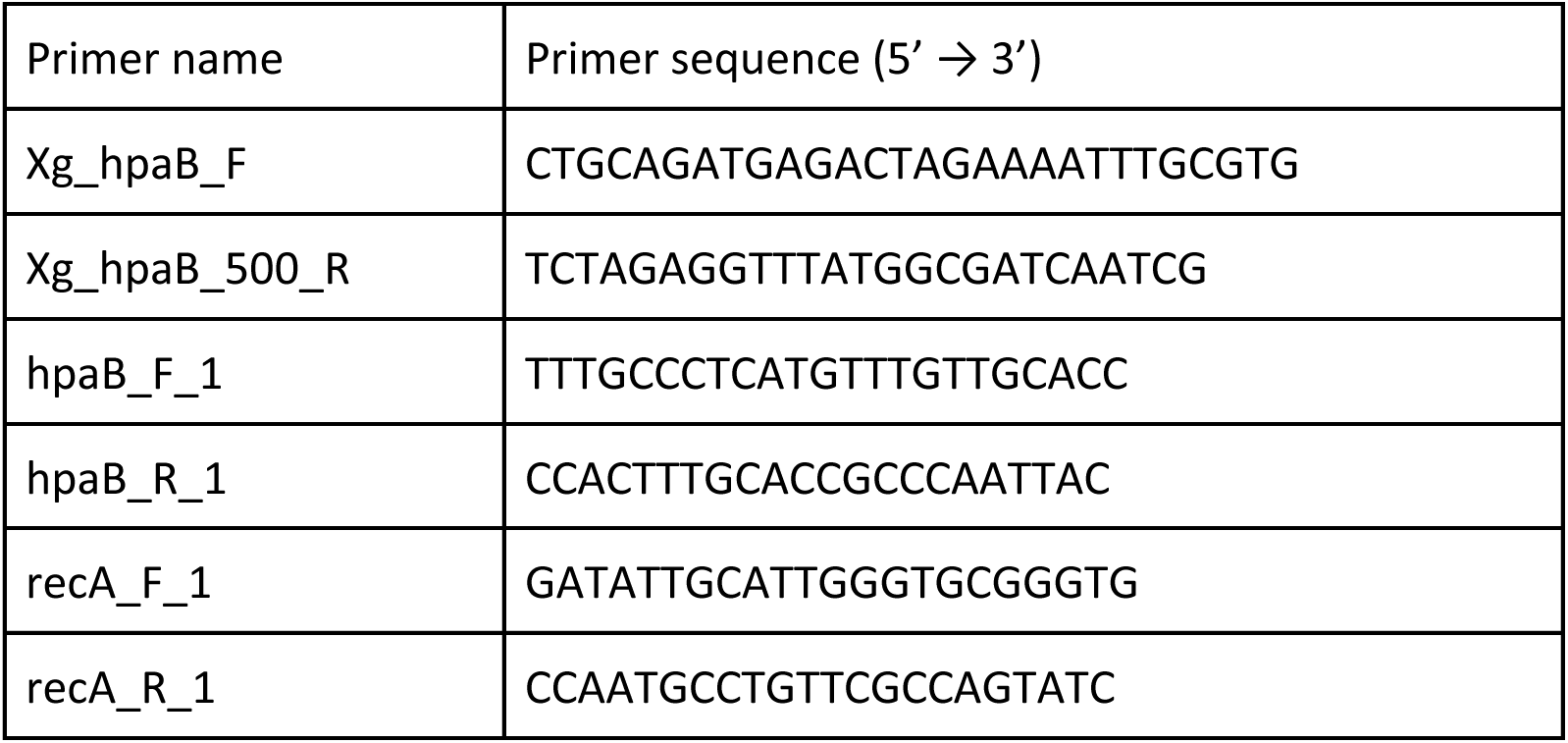
Primers used in this study.

**Table 3.** Bacterial strains and plasmids.

| Strain Name | Genotype | Plasmid | Antibiotic Markers* | Source |
| --- | --- | --- | --- | --- |
| HGB2511 | *X. griffinae* wild type | None | None | Cao et al. (2022) (PMID: 34791196) |
| HGB2622 | *X. griffinae* *P<sub>ara</sub>::hpaBC* | None | Kan | This study |
| HGB2640 | *X. griffinae* *attTn7::GFP* | None | Kan; Chlr | St. Thomas et al. (2024) (PMID: 38371317) |
| HGB2621 | *E. coli* (S17 λpir, DAP-dependent) *P<sub>ara</sub>::hpaBC* donor | pTM003: pCEP_*hpaB_500* | Kan; Requires DAP | This study |
| HGB2383 | *E. coli* (S17 λpir, DAP-dependent) | pCEP_kan | Kan; Requires DAP | Bode et al. (2015) (PMID: 25826784) |

\*Key for abbreviations: Kan = kanamycin, Chlr = chloramphenicol, Strep = streptomycin, DAP = diaminopimelic acid.

### Measuring growth phenotypes of arabinose-inducible strains

To determine whether induction of *hpaBC* expression or the presence of arabinose affected bacterial growth dynamics, we measured the growth of HGB2511 and HGB2622 in liquid culture. Cultures for each biological replicate (*n* = 5) of the bacterial strains analyzed were inoculated from a swipe through of multiple colonies into 5 mL dark LB and incubated overnight at 30°C. The OD_600_ readings of the overnights were used to normalize the cultures which were resuspended and washed with 1 mL of PBS three times. In a 96-well plate, 2.5 µL of washed culture was added to dark LB for a final volume of 250 µL per well. The plate was incubated at 30°C shaking at 280 rpm continuously and the OD_600_ value was read every 15 min for 24 h. The data were plotted using GraphPad Prism v7.05.

### Reverse-transcriptase quantitative PCR (RT-qPCR) of *hpaBC* transcript *hpaB*

To measure arabinose induction of *hpaBC*, liquid cultures of wild-type *X. griffiniae* (HGB2511) and *P_ara_::hpaBC* (HGB2622) were grown overnight at 30°C. These overnight cultures were inoculated into 5 mL dark LB and dark LB + 0.2% arabinose at a 1:50 dilution. After 6 h of incubation (OD_600_ = 1.1-1.4), the equivalent of 1 mL of OD_600_ 1.0 cells was pelleted for each culture, washed in 600 µL Bacterial RNA Protect (Qiagen Cat. No. 76506), and pellets were frozen at -80°C until RNA extraction. RNA was extracted using TRIzol (Invitrogen) and chloroform, followed by precipitation. Briefly, bacterial pellets were resuspended in 50 µL of TE Buffer and 10 µL lysozyme (10 mg/mL). After 3 freeze-thaw cycles, TRIzol and chloroform extraction were performed, followed by isopropanol precipitation and two washes with 70% ethanol. RNA pellets were air dried, resuspended in nuclease-free H_2_O, and concentrations were measured by nanodrop. To generate cDNA, the iScript gDNA Clear cDNA Synthesis kit (Bio-Rad) was used per the manufacturer instructions with 500 ng total RNA input per sample, along with no reverse transcriptase and no template controls. qPCR was performed using 3 µL of a 1:20 dilution of cDNA for each sample, 0.5 µL of 10 mM primers (Table 2) targeting both *hpaB* and *recA* (housekeeping control). Relative fold change of *hpaB* to *recA* transcripts was calculated using the double-delta-CT method, and mean values were compared using a one-way ANOVA with Šídák’s multiple comparison test using GraphPad Prism v7.05.

### *Steinernema* growth assay

Nematode growth in response to bacteria with varying levels of *hpaBC* expression was monitored using nematode length as a proxy. Aposymbiotic eggs and larvae were prepared using an established egg-preparation protocol modified to optimize survival rates for *S. hermaphroditum* (45). Briefly, 2 days after adding IJ nematodes to lipid agar (LA) plates with HGB2511 bacterial lawns (46), adult hermaphrodites were washed off the plates with dH_2_O, treated with 50 mL of bleach solution twice (74 mL of dH₂O, 21 mL of 1.11 M bleach, and 5 mL of 5 M KOH), eggs were pelleted by centrifugation, washed three times with 10 mL dark LB broth, and the washed eggs were incubated overnight at 25°C in dark LB with 25 μg/mL chloramphenicol. The larval concentration was calculated the following day through manual counting, and 1,000 eggs and hatched larvae were added to each of the treatment plates. Treatment plates were prepared by inoculating wild-type and *P_ara_*::*hpaBC* strains (Table 3) in dark LB broth and after incubation overnight at 30°C applying 600 µL of to LA lipid agar (LA) plates, with or without supplementation of 0.2% arabinose. Plates were incubated overnight at 30°C. After incubation at 25°C for 72 h, adult nematodes were washed off plates with 1x phosphate buffer solution (PBS), mounted on microscope slides sealed with VALAP (a 1:1:1 mixture of vasoline, lanolin, and paraffin), and imaged on a brightfield light microscope (Keyence BZ-X710) at 4x and 20x magnification. Nematode lengths were manually traced and measured in Fiji 2.16.0 (47). The lengths of a total of 489 adult hermaphrodites were traced on ImageJ, and any males washed onto the slides were excluded from length-tracing.

### *Steinernema* egg-laying assay

Aposymbiotic *S. hermaphroditum* eggs were generated using the protocol described above. To negate any impacts of *hpaBC* expression on nematode development, nematodes were grown on wild type HGB2511 bacteria until adulthood. Briefly, aposymbiotic eggs and larvae were seeded on lawns of *X. griffiniae* expressing green fluorescent protein (GFP) (HGB2640, Table 3) and incubated at 25°C for 48 h. Nematodes were then washed off the plates with PBS, treated with 100 µg/mL streptomycin for 30 min, pelleted, and deposited onto fresh LB plates with no bacteria to crawl for 2 h. The nematodes were washed off the plates with PBS and distributed onto lipid agar treatment plates (±0.2% arabinose) with wild-type or *P_ara_*::*hpaBC* lawns and incubated overnight at 25°C. A stereo-fluorescence microscope (Leica M165) was used to confirm surface and intestinal clearance of GFP-positive bacteria. Next, 8 hermaphrodites from each treatment were transferred into individual wells of a 96-well plate containing 200 µL PBS and allowed to swim and lay eggs at ambient temperature for 2 h. The number of eggs laid in each well was counted manually under a light microscope.

### Measuring *Xenorhabdus* colonization in IJ nematodes

Aposymbiotic nematode eggs from *S. hermaphroditum* adults were generated and seeded on treatment lawns using the protocol described above. Once these plates were starved of bacteria after 7 d of co-incubation, IJs were collected via White trap and incubated at 25°C for another 7 d to allow for IJ emergence (48). Trapped IJs were then harvested by transferring the water suspension to flasks. Average Colony Forming Units (CFU) per IJ was determined as previously described (49).

### Epifluorescence imaging

Aposymbiotic *S. hermaphroditum* eggs were generated as previously described and placed on lawns of GFP-expressing *X. griffiniae* bacteria (HGB2640) (Table 3). Adult hermaphrodites were picked off the plates using a worm pick, made by embedding a piece of platinum wire into a glass transfer pipette. Nematodes were washed with PBS, immobilized using levamisole, and mounted for imaging using 4% agar pads and imaged using a Keyence BZ-X800 epifluorescence microscope at 20X magnification with brightfield and a GFP filter set.

## Results

### The bacterial HpaB enzyme may bind multiple substrates in the dopamine/tyrosine metabolic pathway based on *in silico* docking predictions

HpaB is the oxygenase subunit of the HpaBC enzyme which catalyzes the hydroxylation of phenolic substrates using O_2_ and electrons from FADH_2_ generated by the HpaC flavin reductase subunit (50,51). In the first step of the homoprotocatechuate pathway, HpaBC converts 4-hydroxyphenol acetate (4-HPA) into homoprotocatechuate (HPC), also known as 3,4-dihydroxyphenylacetic acid (DOPAC), a breakdown product of the neurotransmitter dopamine (Fig. 1) (52). Downstream bacterial homoprotocatechuate pathway enzymes ultimately convert HPC into pyruvate and succinate. HpaBC is present in the genomes of diverse bacterial taxa (53), and our analysis revealed *hpaB* homologs to be widespread in bacterial genomes spanning multiple phyla, including both plant- and animal-associated species, and present on a more limited basis amongst archaea (Fig. S1).

The HpaBC catalytic site may allow potential interactions with phenolic compounds beyond the canonical substrate, 4-HPA (28,29). To explore the idea that microbial HpaBC could potentially interact with animal host-associated molecules such as neurotransmitters and amino acids, we employed a computational docking model to compare docking affinities between the *X. griffiniae* HpaB its canonical substrate, 4-HPA, and two non-canonical substrates, dopamine and L-tyrosine (the shared amino acid precursor of both 4-HPA and dopamine). Docking models derived using SwissDock suggested that the three substrates could adopt similar orientations within the HpaB catalytic site (Fig. 1B). While HpaB is predicted to have higher affinity for 4-HPA over dopamine, association of HpaB with either molecule is predicted to be energetically favorable (Fig. 1C). These findings suggest two plausible mechanisms by which *X. griffiniae hpaBC* expression might alter dopamine metabolism in the host nematode: either by direct hydroxylation of dopamine, L-tyrosine, or other phenolic compounds, or by indirectly impacting flux through the dopamine catabolism pathway through levels of HPC/DOPAC in the host (Fig. 1).

### Generation and validation of an *X. griffiniae* strain expressing arabinose-inducible *hpaBC*

To test whether increasing bacterial *hpaBC* expression has impacts on nematode hosts, we generated an *X. griffiniae* strain with inducible *hpaBC* expression by inserting the arabinose responsive *P_ara_* promoter directly upstream of the *hpaB* gene (Fig. 2A) (42). In this strain, *hpaBC* levels as measured by RT-qPCR, were significantly higher when grown in the presence of arabinose, when compared to the no arabinose added treatment or to the wild-type strain with or without arabinose (Fig. 2B). Neither the addition of arabinose nor the introduction of an inducible promoter system upstream of *hpaBC* significantly impacted the growth dynamics of the *X. griffiniae* strains used in this study (Fig. S2), nor their ability to survive on solid media over the timeframe of the experimental windows used across this investigation (Fig. S3), suggesting that the phenotypes we observe are not due to differential growth or survival of the *X. griffiniae* strains.

**Figure 2.**
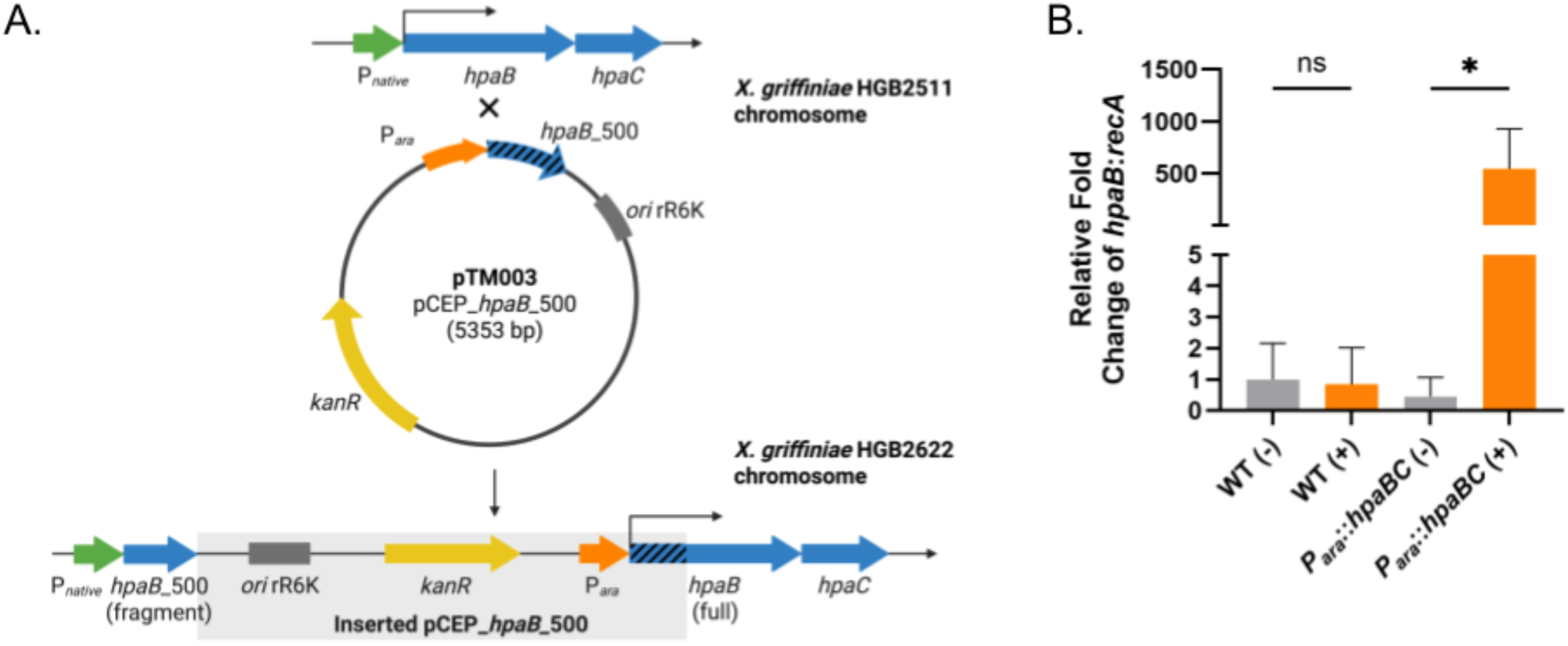
Generation and validation of *X. griffiniae* with arabinose-inducible *hpaBC*. (A) Schematic diagram of the recombination-mediated (X) replacement of the native *hpaBC* promoter (green arrow) with the arabinose-inducible promoter *P_ara_* (orange arrow) upstream of the *hpaBC* genes (blue arrows). The pCEP*_hpaB_*500 plasmid (pTM003) was conjugated into wild-type *X. griffiniae* HGB2511 and selected with kanamycin (kanR, yellow arrow) for single crossover recombination insertion of the plasmid based on *hpaB_*500 homology, generating an arabinose-inducible (HGB2622). (B) RNA was isolated from wild-type (HGB2511) or the inducible *P_ara_*::*hpaBC* strain (HGB2622) grown with (+) or without (-) arabinose. *hpaB* levels were quantified relative to the housekeeping gene (*recA*) by RT-qPCR, and the fold change of *hpaB* relative to *recA* is shown on the y-axis. Significantly higher relative fold change of *hpaB* was observed in the inducible strain *P_ara_*::*hpaBC* in the presence of arabinose relative to the other conditions (*P* < 0.05; one-way ANOVA with multiple comparisons). Error bars depict standard deviation.

### Increased *hpaBC* expression during larval development results in smaller hermaphrodites

Prior to observing impacts on nematode behavior, we assessed whether *hpaB* induction has an impact *Steinernema* nematode development, using hermaphrodite body length as a proxy for rhabditid nematode development (54,55). Adult hermaphrodites grown on *X. griffiniae P_ara_*::*hpaBC* in the presence of arabinose were significantly shorter than those grown on uninduced controls (Fig. 3) (*P* < 0.005). The addition of arabinose alone had no measurable impact on growth, based on the lack of significant differences in length of hermaphrodites grown on wild-type *X. griffiniae* with or without arabinose. This suggests that potential over-expression of *hpaB* (by arabinose induction) in *X. griffiniae* bacteria can negatively impact the growth of the host hermaphrodite if the nematodes are exposed across the juvenile developmental period. The wide variation in body lengths of the adults we observed across conditions was expected because they were not developmentally synchronized.

**Figure 3.**
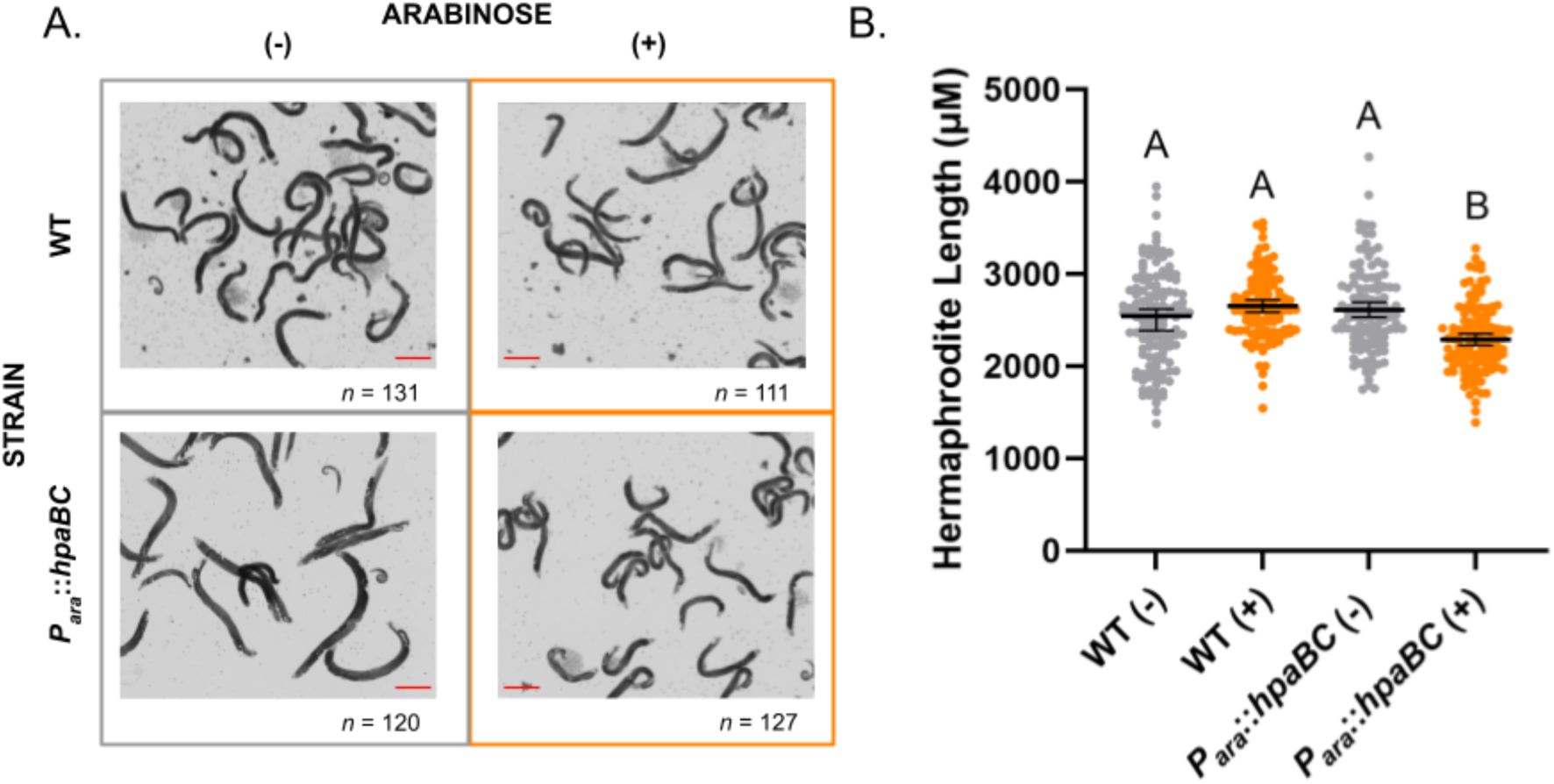
Increased *hpaBC* expression during larval development results in smaller hermaphrodites. (A) Representative images of *S. hermaphroditum* adults across treatment conditions. Aposymbiotic embryos were seeded onto lawns of each indicated strain of *X. griffiniae* grown on lipid agar plates with (+) or without (-) arabinose. Nematodes were allowed to develop into adults, which were washed off the plate and visualized by brightfield microscopy at 4X and 20X. *n* is the total number of hermaphrodites from each treatment whose length was recorded, as shown in B. The scale bar (orange) is 300 μm. (B) The average length of hermaphrodites across each treatment is depicted. Colors reinforce the presence (orange) or absence (grey) of 0.2% arabinose in each treatment. Each point represents a single hermaphrodite. Letters depict distinct statistical groupings as measured by one-way ANOVA (*P* < 0.005), with error bars representing a 95% confidence interval.

### Induction of *hpaBC* expression is inversely correlated with host egg-laying

*Steinernema* adults, including *S. hermaphroditum*, are colonized in the anterior intestine by their *Xenorhabdus* symbionts just posterior to the basal bulb in the anterior intestine (Fig. 4A) (20). However, the impacts of symbiotic bacteria on *Steinernema* adults, including on their behavior, are not well understood. In the free-living nematode *C. elegans,* the availability of nutritive bacteria impacts egg-laying behavior (13,56,57). Moreover, a lack of such bacteria can cause embryos to be retained and hatch within the uterus of the mother. This process, which often results in maternal death, is known as endotokia matricida (58). In *Heterorhabditis* entomopathogenic nematodes, endotokia matricida is thought to be a key step in the vertical transmission of bacterial symbionts from mother to offspring (59). Although a similar transmission process has yet to be documented in *Steinernema* nematodes, a percentage of *Steinernema carpocapsae* mothers do retain eggs which then hatch within the uterus (60). In *S. carpocapsae*, adult colonization of the anterior intestinal cecum by wild-type symbiotic bacteria caused mothers to lay more eggs than nematodes with non-colonizing *rpoS* mutant bacteria (33). Further, the rate of egg-laying in *S. carpocapsae* was also impacted by the exogenous addition of the neurotransmitters serotonin and dopamine, and the predicted product of the HpaBC enzyme overlaps with dopamine catabolism (33) (Fig. 1).

**Figure 4.**
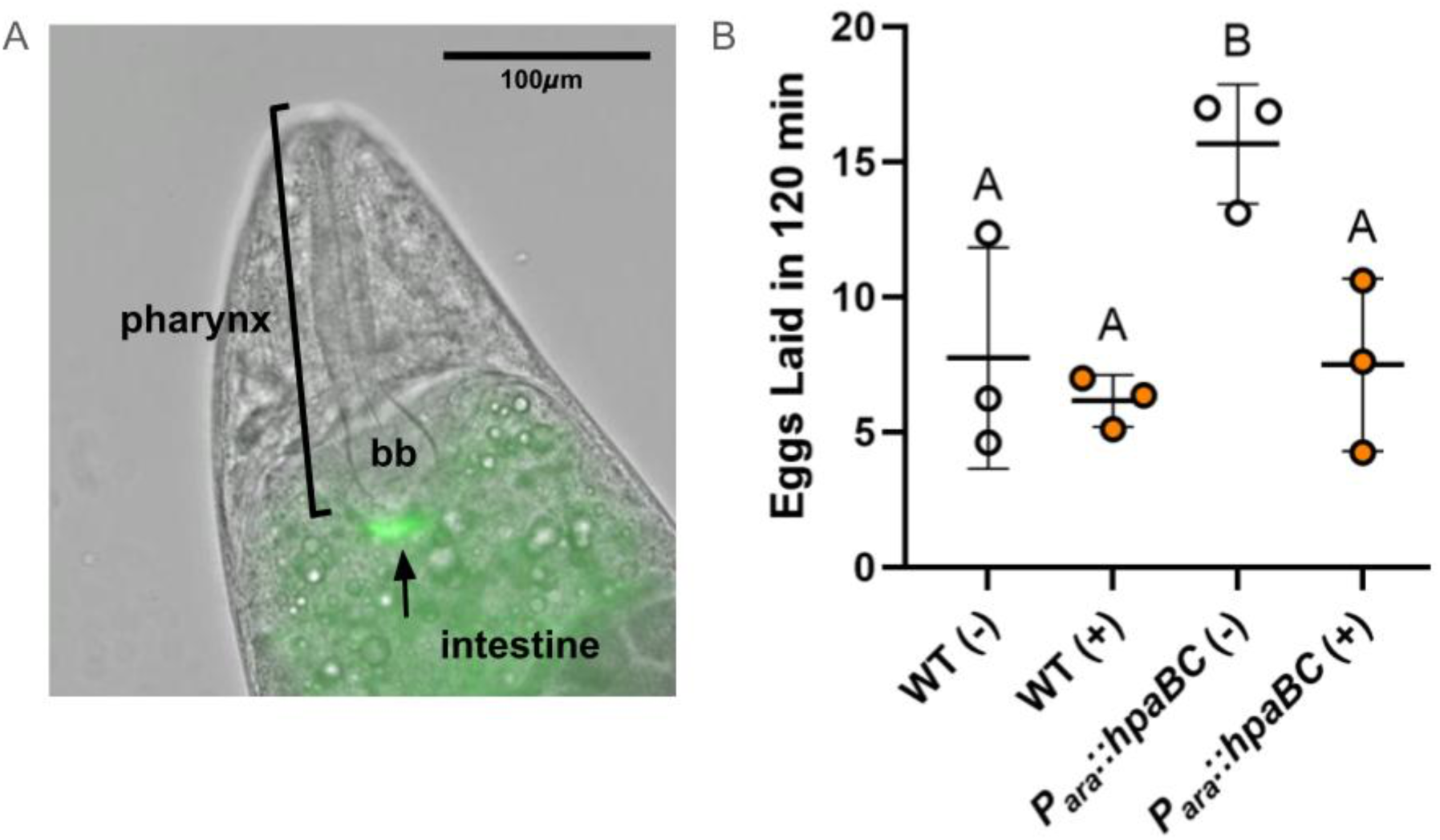
Induction of *hpaBC* expression is inversely correlated with host egg-laying. A) Epifluorescence image showing GFP-expressing *X. griffiniae* bacteria (arrow) colonizing the intestine of an adult *S. hermaphroditum* hermaphrodite just posterior to the basal blub (bb) of the pharynx. B) The mean number of eggs laid by eight hermaphrodites per biological replicate (the three points) over a 120 min period is shown. Colors reinforce the presence (orange) or absence (white) of 0.2% arabinose in each treatment. Letters depict different statistical groupings as measured by one-way ANOVA (*P* < 0.05), with error bars representing standard deviation.

Based on these previous observations, we investigated the impacts of *X. griffiniae hpaBC* levels on *S. hermaphroditum* egg-laying. Because of the impacts of *hpaBC* induction on larval development (Fig. 3), we allowed axenic *S. hermaphroditum* eggs to develop into adults on wild-type *X. griffiniae* bacteria before shifting the hermaphrodites to the treatment conditions for 24 h. Monitoring the egg-laying patterns of the shifted adult hermaphrodites demonstrated that when *X. griffiniae hpaBC* was uninduced, hermaphrodites deposited a significantly higher number of eggs than under conditions of wild-type or arabinose-induced *hpaBC* expression levels (*P* < 0.05) (Fig. 4B). Arabinose-induced over-expression of *hpaBC* did not decrease egg-laying rates below wild-type levels. The presence or absence of arabinose did not significantly influence egg-laying in hermaphrodites treated on wild-type *X. griffiniae* lawns (Fig. 4B). Assuming that, as predicted, the arabinose (-) uninduced condition is similar to an *hpaBC* loss-of-function mutation, these data suggest that the presence of HpaBC suppresses egg-laying in host hermaphrodites through an as yet unidentified mechanism.

### Expression of *hpaBC* is required for *X. griffiniae* persistence within the IJ stage host nematode

As part of their entomopathogenic life cycle, in response to crowding and nutrient deprivation, *Steinernema* nematodes develop into a non-feeding, environmentally-resistant IJ stage that migrates through the soil to seek new insect hosts to infect (61). *Steinernema* nematode hosts are typically colonized with one specific *Xenorhabdus* cognate strain, and the ability of a given *Xenorhabdus* to colonize a nematode host is typically restricted to a single species (62). *Xenorhabdus* is not predicted to be free-living because it cannot produce nicotinamide adenine dinucleotide, a critical vitamin and coenzyme not typically found outside of living cells, and therefore, host colonization is a key fitness determinant for these bacteria (63,64). To test if *hpaBC* expression impacts the ability of *X. griffiniae* to colonize the IJ stage of *S. hermaphroditum*, and therefore be transmitted to subsequent generations, we collected IJs from treatment plates and quantified the average number of bacteria they carried. IJs collected from plates with the uninduced *P_ara_::hpaBC* carried significantly fewer bacteria per IJ nematode than did wild-type and uninduced controls (Fig. 5). Moreover, induction of *hpaBC* expression by the addition of arabinose restores colonization to wild-type levels (Fig. 5). The presence or absence of arabinose alone did not impact wild-type colonization levels, suggesting the difference between the induced and uninduced condition in the *P_ara_*::*hpaBC* strain conditions is the induction of *hpaB* expression. The survival of the bacteria on the plates was robust and not significantly different among the treatment conditions over the time the nematodes were cultured prior to IJ collection (Fig. S3). These results suggest that there is a critical lower threshold of *X. griffiniae hpaBC* expression that is necessary for colonization initiation or persistence within IJs.

**Figure 5.**
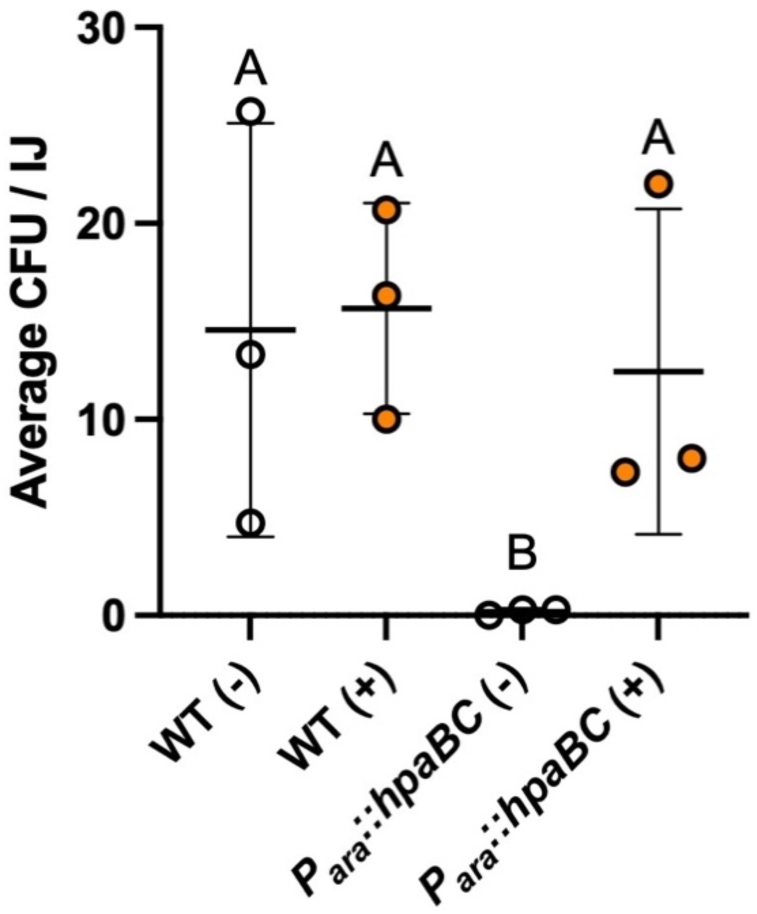
Expression of *hpaBC* is required for *X. griffiniae* persistence within the IJ stage host nematode. Surface sterilized IJs were ground and plated to quantify the average number of colonizing bacteria per infective juvenile (Average CFU/IJ). Points represent biological replicates consisting of the average number of colonies counted across three plates. Colors reinforce the presence (orange) or absence (white) of 0.2% arabinose in each treatment. Letters depict different statistical groupings as measured by one-way ANOVA (*P* < 0.01), with error bars representing standard deviation.

## Discussion

Across animals, evidence is mounting that beneficial host-associated microbes are integrated into the pathways underlying complex traits, including behaviors and development (4). In contrast to the well-established genetic model nematode *C. elegans,* the subject of this study, *Steinernema* nematodes are engaged in a symbiotic partnership with one primary genus of bacteria. Because the pair have evolved together over millions of years of evolutionary time (65), we might reasonably expect the bacterial symbiont to influence aspects of nematode behavior and development, including potentially impacting neuronal signaling pathways. Here, we provide evidence that *Xenorhabdus* can indeed impact nematode development and reproductive behaviors through the expression of *hpaBC*, predicted to encode a tyrosine catabolic pathway monooxygenase.

Using an arabinose-inducible promoter to control *hpaBC* expression, we discovered that over-expression of *hpaBC* results in reduced body length of host nematodes suggesting excess HpaBC may negatively impact larval development (Fig. 3). Though the mechanism of this effect on development is unknown, we verified that it is not due to loss of bacterial viability over the experimental period (Fig. S3). We observed elevated nematode egg-laying in animals cultivated in the uninduced *hpaBC* condition, relative to nematodes cultivated on cells expressing wild-type or elevated *hpaBC* levels, suggesting that the presence of HpaBC negatively modulates egg-laying behavior (Fig. 4). Such retention can lead to endotokia matricida, in which nematode eggs hatch within the body of the mother (66). In some nematodes, endotokia matricida is thought to be a natural part of the lifecycle and a regulated process that optimizes transmission of bacterial symbionts to the next generation (59). Our data suggest that if endotokia matricida is a regulated process in *S. hermaphroditum,* it could be modulated by HpaBC activity, either directly or indirectly (e.g., through action of downstream metabolic products). It is tempting to speculate that via HpaBC, *X. griffiniae* could induce *S. hermaphroditum* endotokia matricida to ensure successful inter-generational transmission from parent to offspring transmission. In a second link to transmission of the bacteria between generations, we found that *hpaBC* induction was necessary for *X. griffiniae* colonization of the IJ nematode (Fig. 5). A requirement for HpaBC in colonization may reflect its involvement in necessary signaling between *X. griffiniae* and the nematode host. Other non-exclusive explanations for the requirement of HpaBC in colonization derive from its potential impacts on bacterial physiology. For example, HpaBC-mediated metabolism might be necessary for survival within the IJ to consume or detoxify metabolites, or HpaBC-derived molecules may be part of a host recognition signaling pathway.

Which HpaB ligands (e.g., 4-HPA or dopamine) and products drive the developmental and behavioral phenotypes we observed remain to be explored. Further, our data do not distinguish between HpaBC or metabolite effects on bacteria versus nematode cell physiology. Our *in silico* docking studies suggest that the catalytic site of *X. griffiniae* HpaB could accommodate either 4-HPA or dopamine in an energetically favorable manner (Fig. 1B). However, we did not identify SecYEG, TAT, or flagellar (Type 3) secretion signals associated with HpaB or HpaC, leading us to predict that the enzyme is localized in the bacterial cytoplasm, rather than being transported to the bacterial periplasm or to nematode host cells (67,68). We found homologs of well-characterized importers for both 4-HPA (HpaX) and tyrosine (TyrP) encoded in the *X. griffiniae* genome (36). Direct functional homologs of the human sodium-dependent dopamine transporter (*SLC6A3*) have not been identified in any bacteria, and the top BLASTP hit in the *X. griffiniae* HGB2511 genome is an uncharacterized sodium-dependent transporter YocR (XGHIN1_v1_3759). Although it is possible that related structural homologs or broad-spectrum amino acid transporters in *Xenorhabdus* could facilitate dopamine uptake, future biochemical assays are necessary to interrogate this possibility. One report in *E. coli* suggests an alternative mechanism for import where dopamine bound to iron enters cells through the enterobactin transport system (69).

A parsimonious model based on currently available information is that HpaBC-mediated metabolism occurs within the *X. griffiniae* cell and that its impact on nematode physiology is indirectly conferred by impacts of 4-HPA metabolites on the bacterium itself or as export products (Fig. 1A). Increased HpaB activity is expected to raise DOPAC levels, as DOPAC is a byproduct of both 4-HPA and dopamine metabolism (Fig. 1A). Because it lacks neurotransmitter function, DOPAC has historically been considered a non-toxic byproduct of dopamine metabolism, however, it is now appreciated that higher levels of DOPAC, in combination with nitric oxide, can lead to mitochondrial dysfunction (70). Elevated levels of DOPAC are often used as a marker for disrupted dopamine metabolism, both clinically and in models of neurodegeneration, including in nematodes (71). Finally, HpaBC could also influence the observed phenotypes through a mechanism unrelated to dopamine signaling or metabolism. An important caveat of our study is that the arabinose-inducible promoter system employed here could be sufficiently leaky to produce *hpaBC* transcript even in the absence of arabinose (72). Indeed, while *hpaB* transcript levels trended lower in the *P_ara_*::*hpaBC* strain under non-inducing conditions relative to wild type, this difference was not significant (Fig. 2). Regardless, the phenotypes we observe here are consistent with HpaBC loss-of-function in the uninduced *P_ara_*::*hpaBC* strain. Testing the impact of the presence, absence, and overexpression of HpaBC on dopamine and other neurotransmitter levels, in addition to the direct impacts of neurotransmitters and their metabolic products on egg-laying behavior, development, and colonization should help clarify these questions.

Taken together, our data show that HpaBC has a negative effect on nematode length and egg-laying but is essential for transmission between generations. Although these effects on diverse developmental and behavioral traits seem disparate at first glance, collectively they suggest that *X. griffiniae* HpaBC activity influences multiple aspects of the decision point between continuing reproduction or exiting to the non-reproductive, insect seeking IJ stage. These results suggest that a metabolic enzyme traditionally associated with aromatic compound degradation can also shape host physiology within an animal-microbe partnership. Because *hpaBC* homologs are widely distributed across bacterial phyla, the ability of bacterial tyrosine metabolism to intersect with host catabolite pathways may represent a broader, novel mechanism through which symbiotic bacteria influence animal biology. This work therefore expands our understanding of how bacterial metabolic activities can become integrated into host developmental and behavioral regulation and provides a framework for exploring similar metabolic interactions across other host–microbe systems.

## Acknowledgements

We thank members of the Goodrich-Blair lab for their help and contributions to this work, especially Erin Mans, for her ground-breaking early work on *X. nematophila hpaBC* and dopamine impacts on *S. carpocapsae.* We thank the Bode lab for their generous gift of the pCEP_kan inducible expression plasmid, the L. Burcham lab for technical support with RT-qPCR, and Dr. Hannah Hughes for providing guidance on computational methods. This work was supported by a grant from the National Science Foundation (IOS-2128266) and the University of Tennessee David and Sandra White Endowment. TGM was supported by awards from the University of Tennessee Advanced Undergraduate Research Activity grant, University of Tennessee Faculty Research Assistants Funding, University of Tennessee Department of Microbiology Summer Research Stipend, and a University of Tennessee Department of Microbiology Jeffrey and Nancy Becker Research Award.

## Author Contributions

Tyler G. Myers: Investigation, Writing - original draft, Visualization

Heidi Goodrich-Blair: Conceptualization, Supervision, Writing - original draft, Writing - review & editing, Funding acquisition

Jennifer K. Heppert: Conceptualization, Investigation, Methodology, Supervision, Writing - original draft, Writing - review & editing, Funding acquisition, Project administration, Visualization

## Supplemental Figures

**Supplemental Fig. 1:**
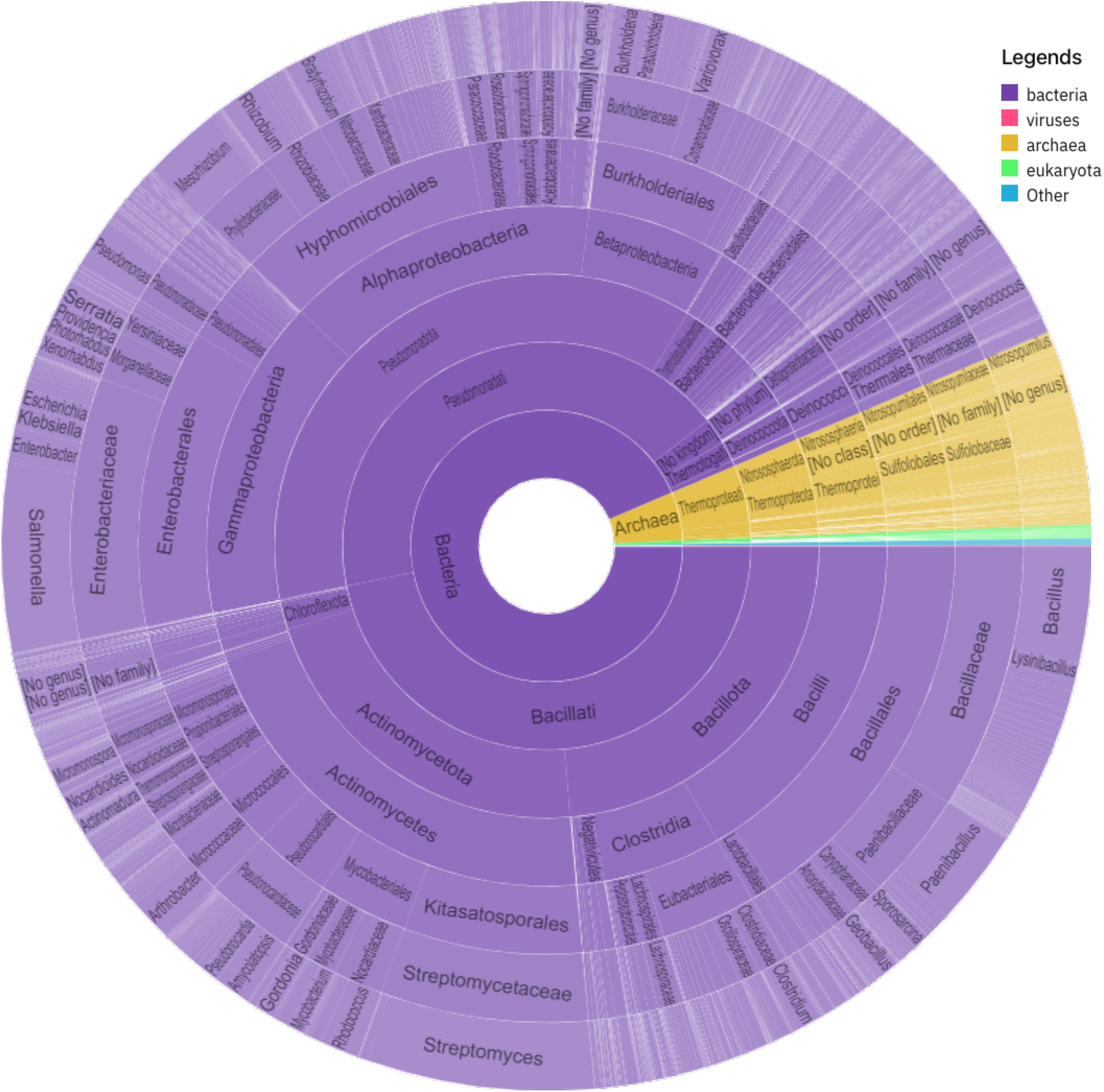
Taxonomic distribution of proteins containing the HpaB/PvcC/4-hydroxyphenylacetate monooxygenase domain. Taxonomic distribution of HpaB homologs was assessed using the InterPro entry IPR004925 (HpaB/PvcC/4-BUDH family) within InterPro v106.0 (https://www.ebi.ac.uk/interpro/entry/InterPro/IPR004925/taxonomy/uniprot/#sunburst). InterPro aggregates protein-family signatures from multiple member databases and maps matching UniProtKB sequences to the NCBI taxonomy tree. The InterPro taxonomy viewer was used to generate a sunburst plot summarizing the number of species containing proteins annotated with this domain, which was exported directly from the InterPro interface. Widths of taxa are weighted by the number of species contained. Color represents the domain to which each HpaB homolog species belongs, as indicated in the legend.

**Supplemental Fig. 2:**
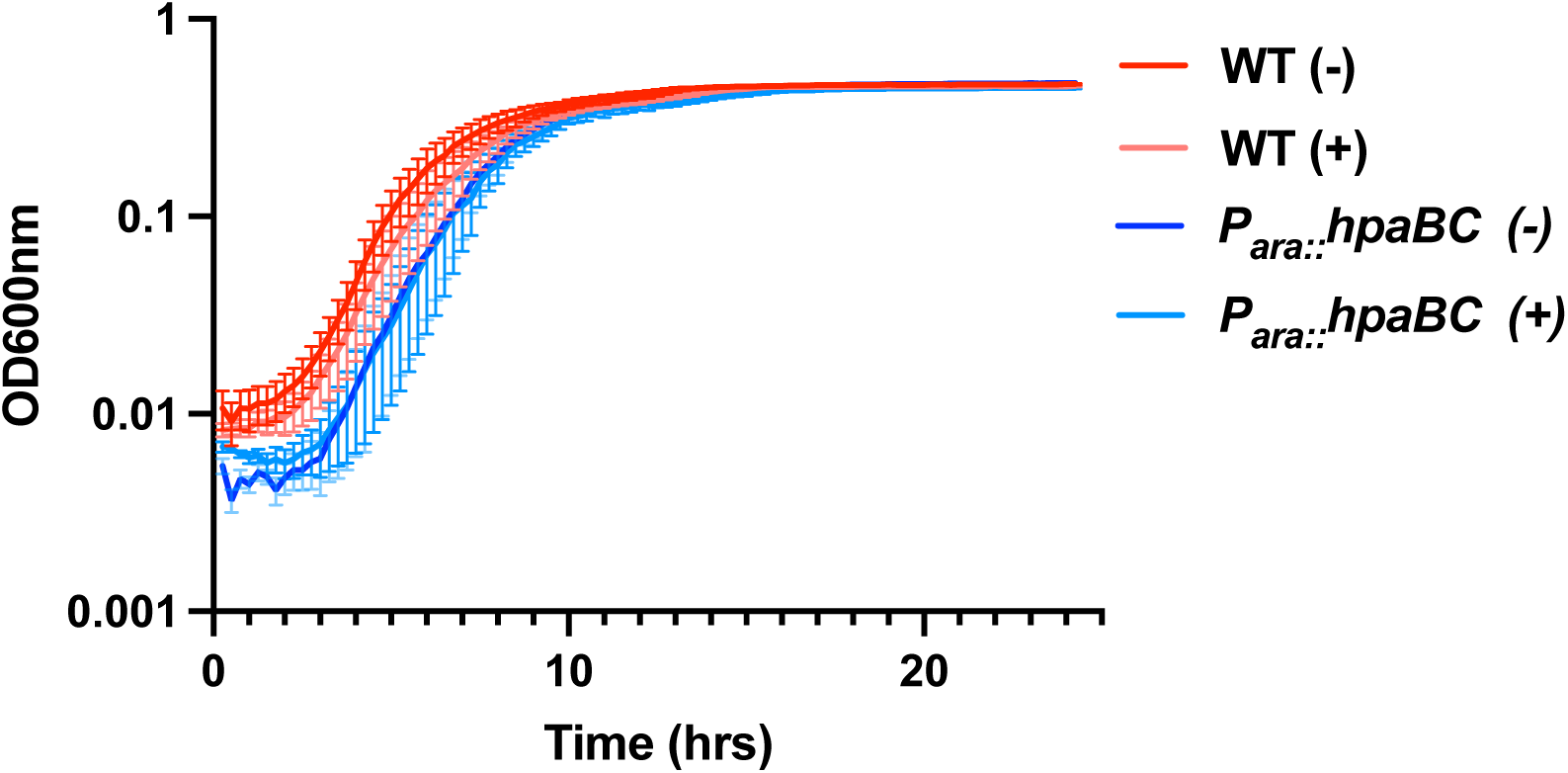
Growth in liquid culture of wild-type and *P_ara_*::*hpaBC* . Growth curves for 4 biological replicates of each strain were performed by measuring OD_600_ over 25 hours. Error bars represent Standard Error (SEM). The curves were fit using a least squares method in Graph Pad PRISM (Exponential growth with log(population)). The doubling rate for each curve was determined and the mean for each condition was compared using an Ordinary one-way ANOVA with multiple comparisons. The doubling rates were found to be non-significantly different among the conditions. An estimate of the carrying capacity was compared using an average maximum OD for each curve in stationary phase (the 20-25 h time points). An Ordinary one-way ANOVA with multiple comparisons revealed that the P*_ara_*::*hpaBC* strain with arabinose OD during stationary phase was significantly different from the P*_ara_*::*hpaBC* strain without arabinose (p= 0.0067). However, it was not significantly different from the wild type strain with arabinose (p= 0.4460), suggesting that the effect may be due in part to the presence of arabinose. The difference was small, with the average maximum OD for all curves ranging from 0.432-0.477.

**Supplemental Fig. 3:**
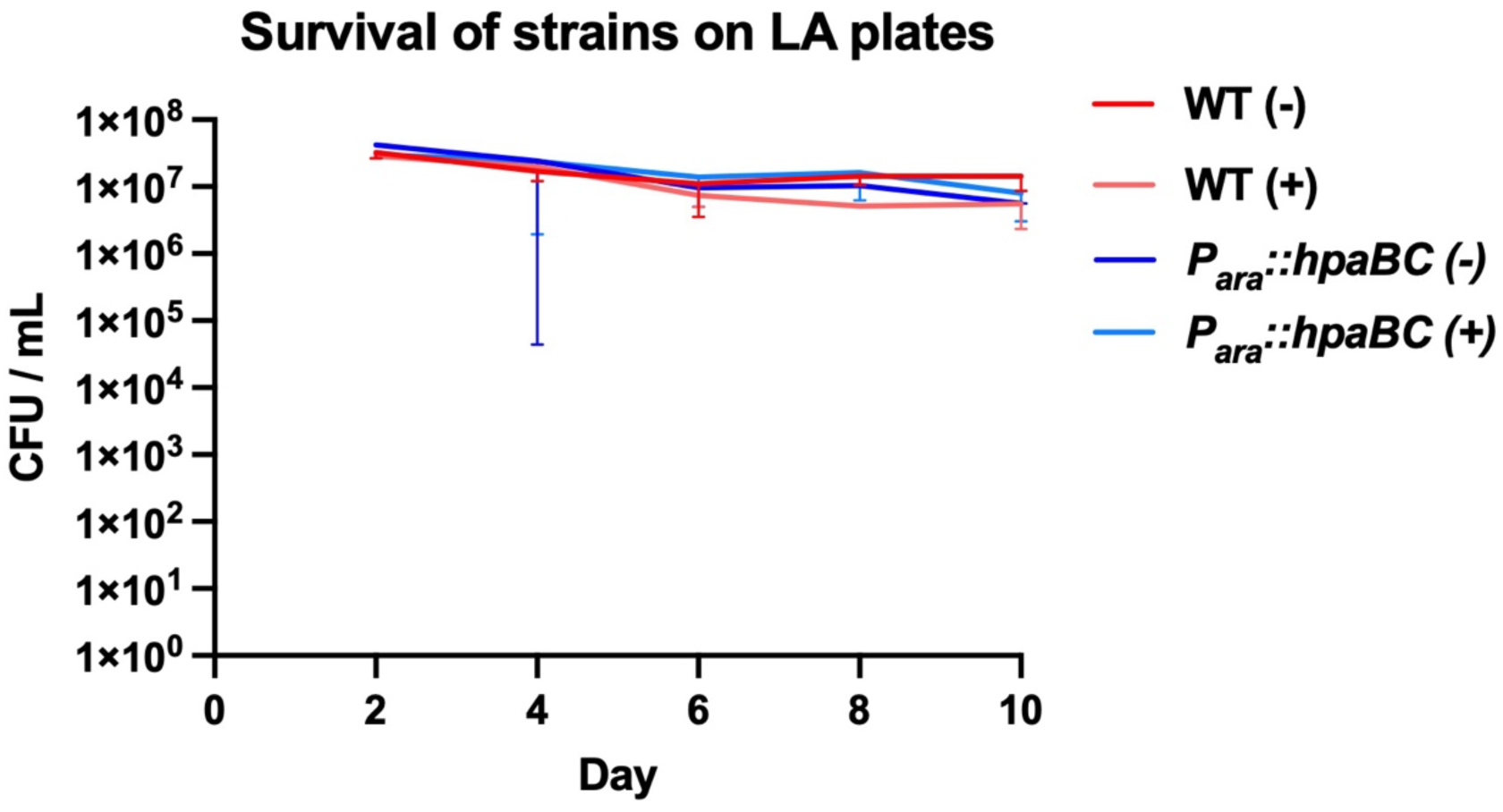
Survival on agar lawns of wild-type and *P_ara_*::*hpaBC*. The survival of wild type (WT) and *P_ara_::hpaBC* strains on lipid agar plates with (+) and without (-) arabinose was assessed over the course of 10 d. CFU/ml were counted each day. Error bars represent 95% confidence intervals. A Two-way ANOVA with repeated measures was used to determine whether day, replicate, or condition were significantly different across the measurements. The decrease in CFU over the days was significant (p=0.0017), but there was no significant difference between replicates (p=0.6441) or conditions (p=0.2480).

**Supplemental Table 1 (Excel file): Tyrosine and Dopamine Catabolism in *X. griffiniae* and *S. hermaphroditum.*** Excel sheet with the *X. griffiniae, S. hermaphroditum,* and *C. elegans* homologs of enzymes in the tyrosine and dopamine catabolism pathways identified in this study and named as labels for the reactions in Figure 1. Enzyme name, reaction catalyzed, KEGG ID, locus tag, and amino acid sequences are given. The * symbol denotes genes in the *C. elegans* genome with paralogs that may carry out a similar function as the gene listed in the table. The # symbol denotes that the *comt-4* gene we chose is one of 5 homologs of mammalian COMT present in the *C. elegans* genome. We chose *comt-4* as a homolog for comparison, as behavioral evidence suggests linkage as most closely associated with dopamine metabolism (73).

